# A single-molecule nanopore sequencing platform

**DOI:** 10.1101/2024.08.19.608720

**Authors:** Jia-Yuan Zhang, Yuning Zhang, Lele Wang, Fei Guo, Quanxin Yun, Tao Zeng, Xu Yan, Lei Yu, Lei Cheng, Wei Wu, Xiao Shi, Junyi Chen, Yuhui Sun, Jingnan Yang, Rongrong Guo, Xianda Zhang, Liu’er Kong, Zong’an Wang, Junlei Yao, Yangsheng Tan, Liuxin Shi, Zhentao Zhao, Zhongwang Feng, Xiaopeng Yu, Chuang Li, Wu Zhan, Yulin Ren, Fan Yang, Zhenjun Liu, Guangnan Fan, Weilian Zhong, Dachang Li, Lei He, Yanwei Qi, Meng Zhang, Yening Zhu, Heng Chi, Ziyu Zhao, Zhuofang Wei, Ziqi Song, Yanmei Ju, Ruijin Guo, Liang Xiao, Xiumei Lin, Liang Chen, Chentao Yang, Qiye Li, Ou Wang, Xin Jin, Ming Ni, Wenwei Zhang, Longqi Liu, Ying Gu, Jian Wang, Yuxiang Li, Xun Xu, Yuliang Dong

## Abstract

Nanopore sequencing, a third-generation sequencing technology, has revolutionized the gene sequencing industry with its advantages of long reads, fast speed, real-time sequencing and analysis, and potential in detecting base modifications. This technology allows researchers to sequence longer DNA fragments in a single read, providing more comprehensive genomic information compared to previous methods. Nanopore sequencing operates on electrical signals generated by a nanopore embedded in a membrane separating two electrolyte-filled chambers. When single-stranded DNA (ssDNA) passes through the nanopore, it creates variations in the current that correspond to different DNA bases. By analyzing these current fluctuations with machine learning algorithms, the DNA sequence can be determined. In this study, we introduced several improvements to nanopore sequencing, including nanopore local chemistry sequencing, novel motor and pore proteins, chip design, and basecalling algorithms. Our new nanopore sequencing platform, CycloneSEQ, demonstrated long-duration sequencing (107 hours) on a single chip with high yield (>50 Gb). In human genomic DNA sequencing, CycloneSEQ was able to produce long reads with N50 33.6 kb and modal identity 97.0%. Preliminary findings on human whole-genome *de novo* assembly, variant calling, metagenomics sequencing, and single-cell RNA sequencing have further highlighted CycloneSEQ’s potential across different areas of genomics.

## 1 Introduction

Nanopore sequencing, which has emerged as a novel sequencing technology in recent years, has revolutionized the gene sequencing industry due to its advantages of long reads, real-time sequencing, portability, and minimal library preparation [1]. This technology enables researchers to sequence longer fragments of DNA in a single read, providing more comprehensive genomic information compared to previous methods. Nanopore sequencing is based on electrical signals [2]. It involves a nanopore, which can be either a protein or solid-state structure, embedded in a membrane that separates two electrolyte-filled chambers. When a voltage is applied across these chambers, it generates a steady transmembrane current. As molecules, such as single-stranded DNA (ssDNA), enter the nanopore, they obstruct the flow of ions, creating variations in the current known as nanopore signals. The obstruction of the current varies with different DNA bases (adenine, thymine, cytosine, and guanine) as the ssDNA passes through the nanopore. By detecting these current fluctuations and analyzing them using machine learning algorithms, the DNA sequence can be determined [3] [4] [2].

The development of nanopore sequencing can be traced back to the 1980s, encompassing stages of concept validation, technology development, and commercial application. In 1989, scientists George Church, David Deamer and Daniel Branton proposed the concept of using nanopores for DNA sequencing [5]. In 1996, Kasianowicz and colleagues first demonstrated the phenomenon of current blockade by DNA molecules passing through an *α*-hemolysin protein nanopore, laying the foundation for nanopore sequencing [6]. In 1997, Deamer and Akeson further validated that the current signals produced by single nucleotides passing through a nanopore could be used to distinguish different nucleotides [7]. In 2012, Oxford Nanopore Technologies launched the first portable nanopore sequencer, the MinION, marking the entry of nanopore sequencing into practical application. In 2016, Oxford Nanopore Technologies introduced the PromethION, a high-throughput nanopore sequencing platform that further enhanced sequencing speed and accuracy. Currently, nanopore sequencing technology is widely applied in genomics, transcriptomics, epigenetics, and clinical diagnostics [5].

In this study, we experimented with several improvements to the nanopore sequencing technology, including novel motor and pore proteins, chip design, basecalling algorithms, and nanopore local chemistry (NLC) sequencing method. We demonstrate our new nanopore sequencing platform, CycloneSEQ, is able to perform long-duration (107 h) sequencing on a single chip with high yield (>50 Gb). Whole-genome sequencing of the HG002 cell line produced long reads with N50 33.6 kb and modal identity 97.0%. Further data analyses of CycloneSEQ confirmed its capability of high-throughput long-read sequencing and potential in genomic, metagenomic and epigenomic applications. Preliminary down-stream analyses on human whole-genome *de novo* assembly, variant calling, metagenomics sequencing and single-cell RNA sequencing further demonstrated the potential of CycloneSEQ in various domains of genomics.

## 2 Results

### 2.1 Screening of motor and pore proteins

Motor proteins and pore proteins are two key components of a nanopore sequencing system, playing crucial roles in the precise and efficient sequencing of nucleic acids [8]. We selected helicases as our motor proteins for nanopore sequencing due to their inherent ability to unwind dsDNA, a critical function for sequencing applications. Through comprehensive sequence and structural searches within deep-sea metagenomic databases, we identified numerous motor proteins with novel sequences and structures. These newly discovered proteins exhibit low sequence homology (approximately 35%) to known helicases, indicating their unique evolutionary paths and potential for novel functionality. The structures of these proteins and ssDNA complexes predicted by AlphaFold3 [9]show that they possess distinct helicase characteristics and exhibit significant structural novelty compared to known structures (Fig. 1a).

**Figure 1.**
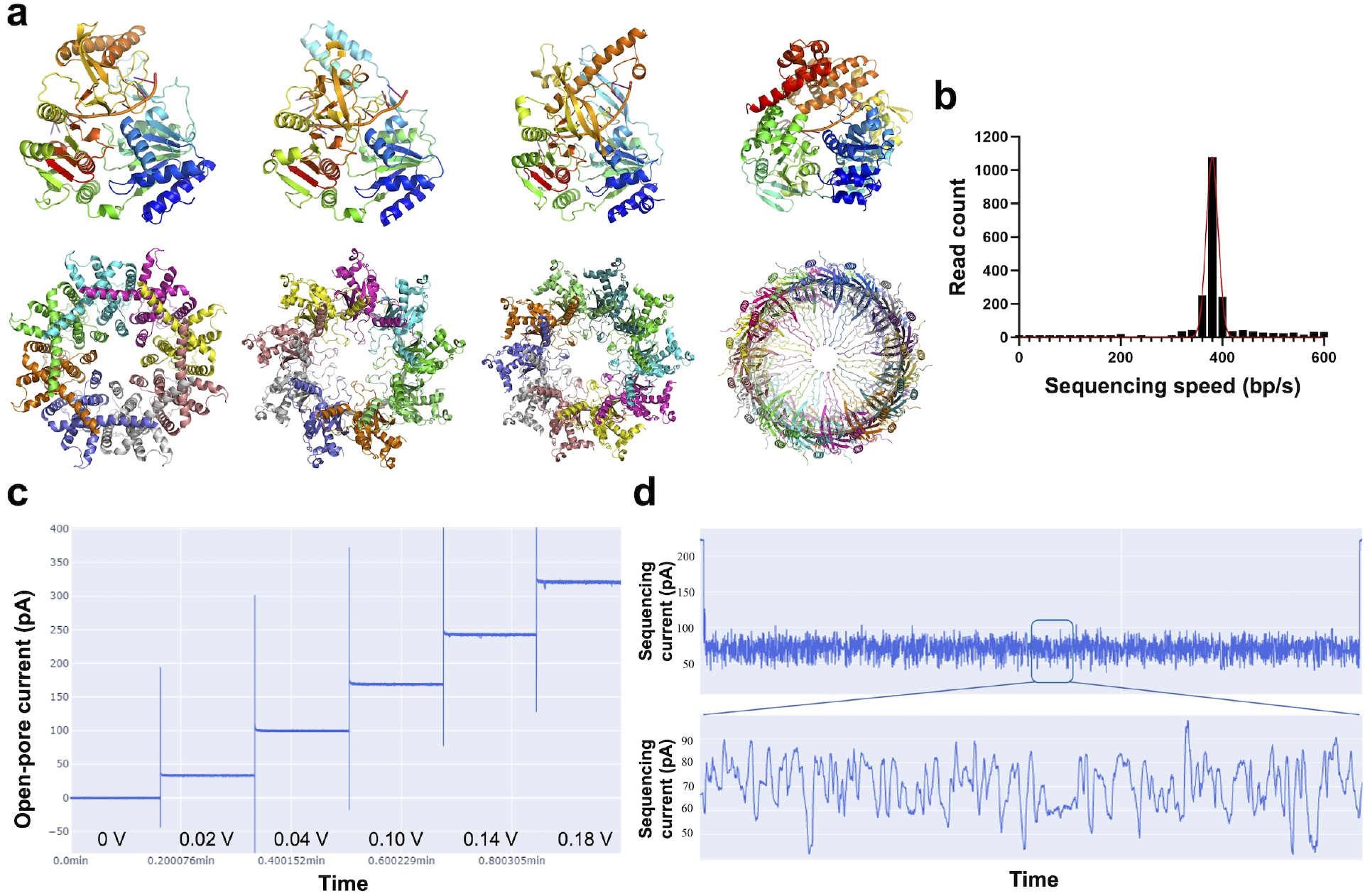
Screening of motor and pore proteins. **(a)** Schematic diagram of AlphaFold3 structure prediction of candidate helicases (top) and pore (bottom) proteins. **(b)** Distribution of nanopore sequencing speed of helicase BCH-X. **(c)** Voltage ramping study of pore portein BCP-Y, with voltages set at 0 V, 0.02 V, 0.04 V, 0.10 V, 0.14V and 0.18 V. **(d)** Representative nanopore sequencing current signal of a single DNA strand generated by helicase BCH-X coupled with pore protein BCP-Y. The magnified section displays “current squiggles” caused by different nucleotides translocating through the nanopore.

Following extensive experimental screening and mutational engineering, we found that most of these motor proteins were well-suited for nanopore sequencing. For example, BCH-X, a member of candidate proteins, demonstrated strong DNA binding and 5’ to 3’ DNA unwinding activity. This activity is essential for maintaining the progression of DNA strands through the nanopore. By screening mutants of BCH-X, we achieved a sequencing speed of approximately 380 bp/s under our sequencing conditions, with high uniformity (Fig. 1b). High speed uniformity is critical in nanopore sequencing because it ensures consistent data output and reduces the likelihood of errors, thereby enhancing the overall accuracy and reliability of the sequencing process.

In tandem with motor proteins, pore proteins are integral to the function of nanopore sequencing systems, as they form the channels through which nucleic acids are translocated and detected. Similarly to our approach with motor proteins, we identified several different families of pore proteins with novel sequences and structures from deep-sea metagenomic databases. These proteins exhibit less than 50% sequence homology to known pore proteins, highlighting their potential for providing new insights and capabilities in sequencing technologies. AlphaFold3[9] structure prediction and protein preparation results show that they can form a nanoscale channel structure as a homomultimer. (Fig. 1a) Using BCP-Y as an example, in pore insertion experiments, BCP-Y can efficiently embed into the membrane and exhibits low-noise open pore currents at different voltages, demonstrating its potential for application in nanopore sequencing (Fig. 1c). By screening a large number of mutants (especially in the sensor region) and combining them with the motor protein BCH-X, the pore protein BCP-Y can facilitate ssDNA translocation and sequencing with high signal complexity and good signal-to-noise ratio of the sequencing current signal (Fig. 1d). This ultimately led to a significant improvement in the accuracy of BCP-Y nanopore sequencing. Additionally, novel structural features at the BCP-Y “lip” motif (abundant positive charges) endow it with enhanced nucleic acid capture capabilities, facilitating more efficient DNA threading through the pore and thereby contributing to the overall efficiency and reliability of the sequencing process.

### 2.2 Pre-training and fine-tuning of the basecalling algorithm

Existing basecalling models for nanopore sequencing primarily use supervised training, requiring large amounts of labeled sequencing data [10]. This method is costly and involves extensive training cycles. When data is insufficient, prediction accuracy suffers, leading to high costs and low accuracy [8]. We adopted a pre-training and fine-tuning approach to address these issues (Fig. 11). During pre-training, the model learns from vast amounts of unlabeled data, allowing it to “understand” the data. Fine-tuning then uses pre-trained weights for rapid convergence and enhanced accuracy. Inspired by Facebook’s wav2vec 2.0 [11], a pre-training method for speech tasks, we employed it for basecalling. Wav2vec 2.0 uses a large corpus of unlabeled speech data for pre-training and a small amount of labeled data for fine-tuning downstream tasks, demonstrating this approach’s feasibility.

Pre-training on a diverse dataset covering various species reduces error rates and accelerates convergence. Future directions include scaling up the model and training samples for enhanced accuracy and expanding species diversity in training data for broader downstream task support. Additionally, leveraging weak label data, as demonstrated by OpenAI’s Whisper model, could enhance sequencing models’ robustness and utility.

### 2.3 Nanopore local chemistry (NLC): a novel method for single-molecule sequencing

The local chemical environment within or near the nanopore is a critical determinant of the performance and accuracy of nanopore sequencing technologies. Variations in local ion concentration, pH, and the presence of other molecular species can significantly influence the ionic current, biochemical reactions, and consequently, the detection and discrimination of nucleotides as they translocate through the pore. The local ion concentration, in particular, affects the electrostatic landscape of the nanopore, which can alter the speed and behavior of nucleic acid molecules during sequencing. Understanding and controlling the local chemical conditions are therefore essential for optimizing the sequencing process, reducing error rates, and achieving high-fidelity reads. Ongoing research into the local chemistry of nanopores aims to elucidate the complex interplay between these factors and to develop strategies for maintaining optimal conditions throughout the sequencing run.

Beyond optimizing data quality by manipulating the local chemical environment near the pore, we introduced a novel sequencing method, termed nanopore local chemistry (NLC) sequencing. We first created an asymmetric chemical environment on each side of the nanopore. On the *cis* side, the sequencing buffer contained no magnesium ions (Mg^2+^), while the electrolyte on the *trans* side contained 20 mM magnesium ions (Fig. 2a). DNA helicase requires both magnesium ions and ATP to properly unwind the DNA double helix. Specifically, magnesium ions first bind with ATP to form an Mg-ATP complex. This complex, which is the actual substrate for DNA helicase, can be recognized and utilized by the helicase. When we introduced library molecules (a mixture of dsDNA and helicase) on the *cis* side, the DNA double helix could not be properly unwound due to the lack of Mg^2+^ (Fig. 2a).

**Figure 2.**
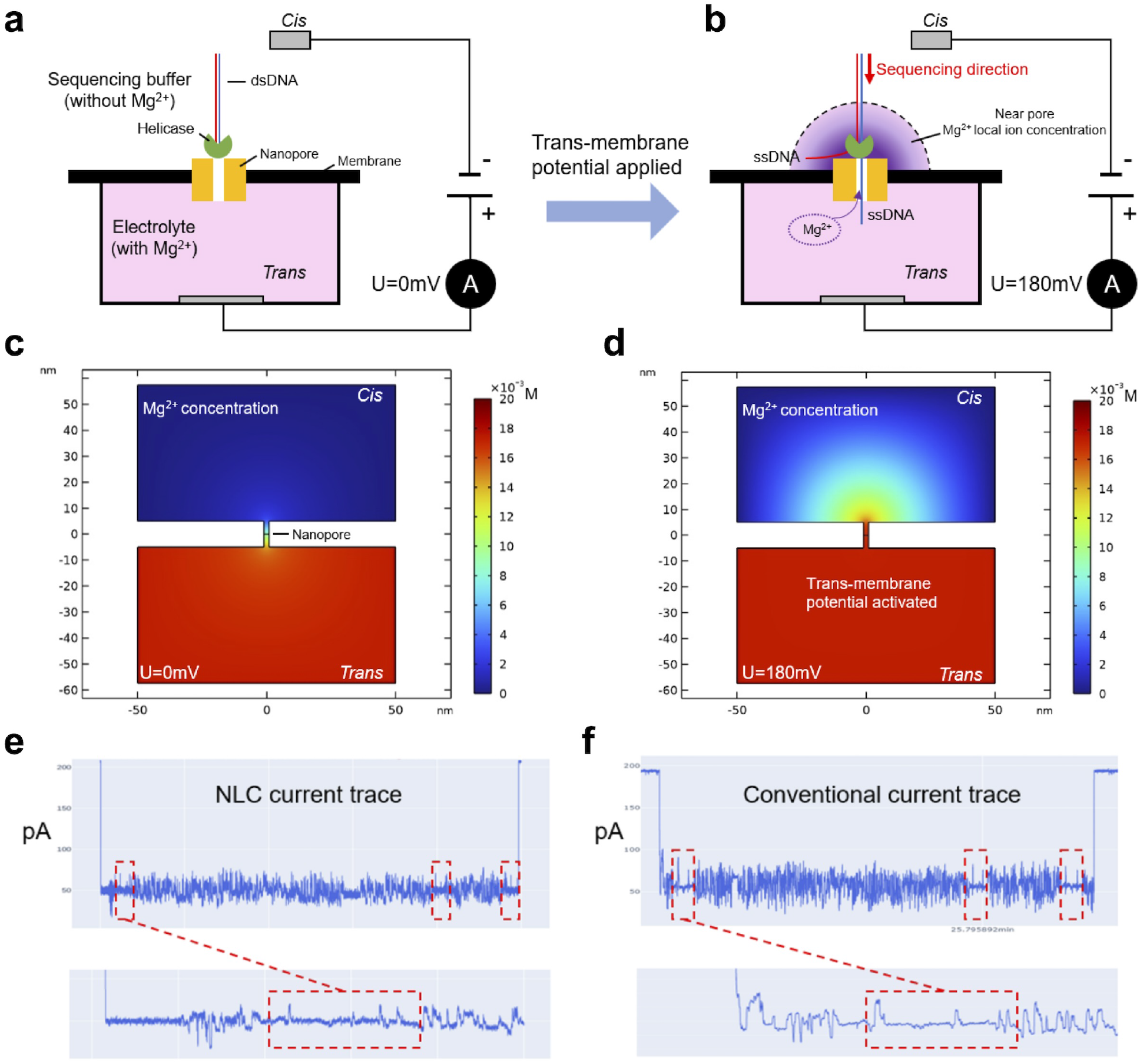
The nanopore local chemistry (NLC) sequencing method. **(a)** Schematic representation with no trans-membrane potential applied (*U* = 0 mV). Magnesium ions are present in the *trans* microwell, but not in the *cis* microwell, preventing dsDNA unwinding. **(b)** When the trans-membrane potential is activated (*U* = 180 mV), magnesium ions translocate from the *trans* microwell to the *cis* side through the nanopore, creating a localized concentration of magnesium ions. **(c)** COMSOL simulation of Mg^2+^ ion concentration distribution (*U* = 0 mV). **(d)** COMSOL simulation of Mg^2+^ ion concentration distribution when a trans-membrane potential is activated (*U* = 180 mV). **(e-f)** Sequencing current trace of nanopore local chemistry (NLC) method **(e)** and the conventional **(f)** method. The library DNA molecules contain three repetitive sequences, which are successfully reflected in the current signals of both sequencing methods (indicated by red dashed boxes).

However, after applying the transmembrane potential (*U* = 180 mV), Mg^2+^ ions were transported from the *trans* microwell to the *cis* side through the nanopore, creating a local concentration gradient of Mg^2+^ on the *cis* side near the pore (Fig. 2b). According to the simulation results (Fig. 2c), the Mg^2+^ concentration maximized near the pore and decayed rapidly. Library molecules captured by the nanopore electric field were pulled near the nanopore entrance and exposed to the Mg^2+^-rich environment. Magnesium ions near the pore bound with ATP to form an Mg-ATP complex, which was then utilized by the helicase, thus activating sequencing (Fig. 2d).

The ionic current trace produced by NLC sequencing are shown in Fig. 2e compared with conventional nanopore sequencing current trace (Fig. 2f). In both cases, we applied a transmembrane potential of 180mV. The open pore current, when there isn’t DNA translocating through the pore, of NLC is 206.70 pA with and a standard deviation of 0.51 pA. The open pore current of conventional nanopore sequencing is 193.91 pA with a standard deviation of 0.76 pA, respectively. The mean current during sequencing, when DNA passing through the pore, for NLC is 49.1 pA, with a standard deviation of 7.25 pA, an amplitude of 61.22pA. The mean current during sequencing for conventional nanopore sequencing method is 57.17 pA, with a standard deviation of 11.00 pA, an amplitude of 68.05pA. To draw a conclusion, both methods exhibit very similar characteristic current values. As shown in Fig. 2 e-f, the library DNA molecule (approximately 1.5 kb in total length) contains three repetitive sequences, which are successfully reflected in the current signals of both sequencing methods.

### 2.4 A nanopore sequencing platform based on improved chip design

The design of the nanopore sequencing chip is a critical factor in the advancement of nanopore sequencing technology, which offers a unique approach to DNA and RNA sequencing by monitoring changes in ionic current as nucleic acids pass through a biological nanopore. This technology relies on a sensor chip designed with arrays of microwells, which support membrane arrays and contain microelectrodes at the bottom of each well. In this setup, biological nanopores are inserted into the membrane arrays that are uniformly formed on the sensor chip. The membranes are self-assembled in a bilayer form via lipid molecules. Each nanopore is electrically connected to electrodes that precisely measure the ionic current disruptions caused by nucleotide sequences moving through the pore.

The core metrics of a nanopore sequencer are primarily sequencing throughput and accuracy. To enhance sequencing throughput, we employed high-density nanopore arrays on the sequencing chip and optimized the spatial distance between nanopores to maximize the parallel processing of nucleic acid strands. The pitch distance between each microwell is around 200 µm, resulting in a maximum nanopore density of approximately 28.9 per mm^2^(Fig. 3a).

**Figure 3.**
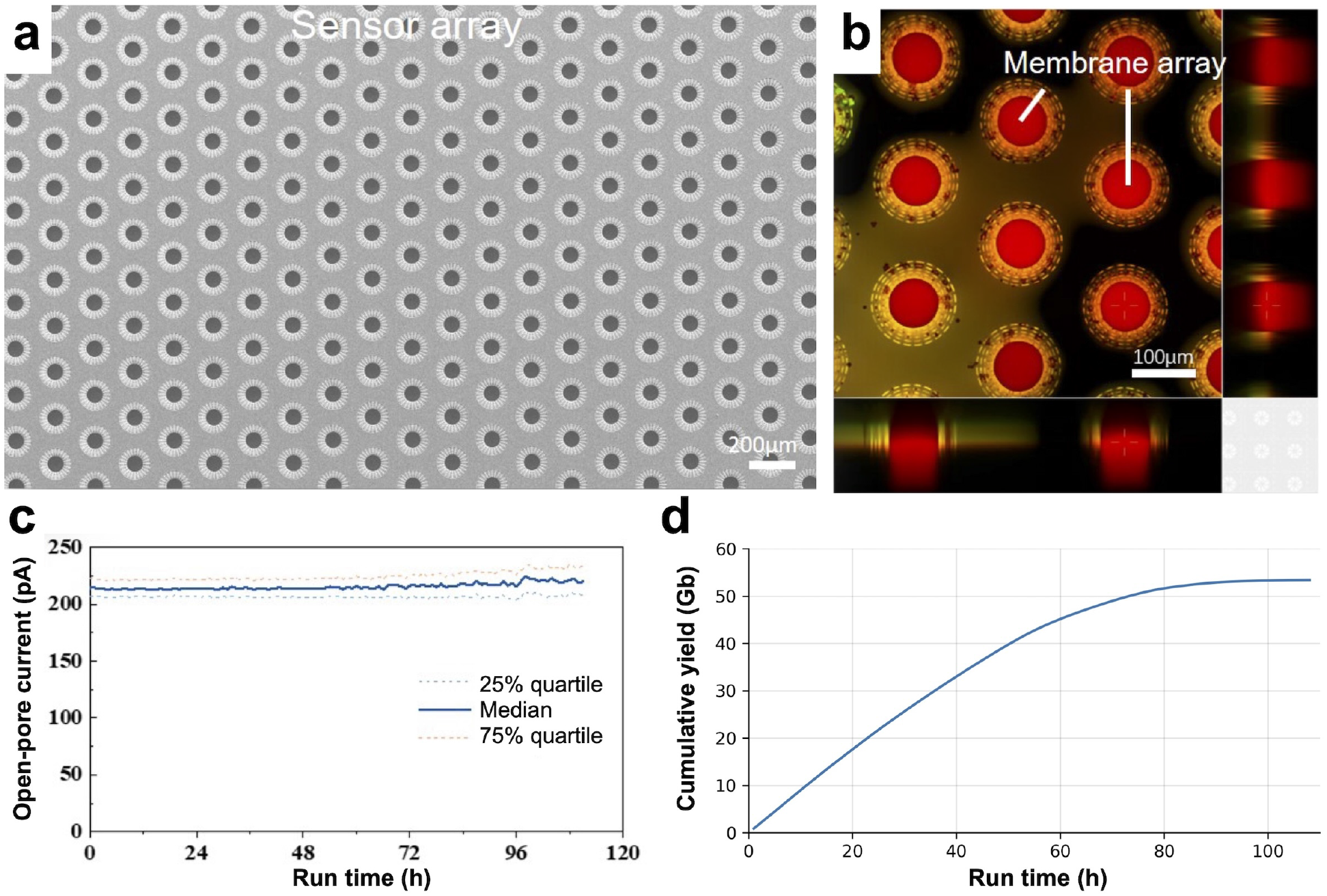
Improved chip design for nanopore sequencing. **(a)** Scanning electron microscope (SEM) micrograph of sensor arrays for nanopore sequencing. Sensor units are spaced approximately 200 µm apart and are arranged in a honeycomb pattern to maximize sensor density. **(b)** Confocal image of the membrane array formed on the sensor chip. Electrolyte in each microwell is shown in red, while the membrane solution is shown in green. **(c)** Median open pore current of all effective single nanopores on a sensor chip during a 112-hour sequencing run. **(d)** Cumulative number of bases sequenced over time in a 107-hour sequencing on a single flowcell.

Additionally, we engineered the microwell wall structure to maximize the electrolyte buffer volume within each well, leading to an electrochemical system with prolonged stability.

To improve sequencing accuracy, we focused on enhancing the signal-to-noise ratio of the chip. This involves implementing smaller apertures for the microwells. In our system, minimizing the size of the aperture (with a diameter of ≤76 µm) results in a smaller final membrane area (Fig. 3b). A smaller membrane area leads to lower membrane capacitance (≤20 pF) and reduced noise, which is electrically coupled to the measuring system.

These design improvements enabled our sensor chip to support over four days of continuous sequencing with consistent open-pore currents (Fig. 3c). We sequenced the *E. coli* genome for 107 hours on a single flowcell. This sequencing run cumulatively yielded 53.4 Gb data that passed the internal basecalling quality criteria (Fig. 3d), demonstrating the possibility to achieve high sequence yield by sequencing for a prolonged period of time. Among the 12.6 million reads generated, 12.1 million (95.6%) was mapped to the *E. coli* reference genome. We note that no buffer re-flush or library washing was employed here, which are common ways to maintain sequencing speed and accuracy.

Based on the novel nanopore sensor chip design described above, we have successfully constructed a nanopore based single-molecule sequencing platform named as CycloneSEQ. As illustrated in Fig. 4b, The flow cell module of CycloneSEQ comprises a microfluidic chip enabling the transportation and temporary storage of sample molecules as well as supporting electrochemical reaction, an arrayed chip containing nanopores, a signal acquisition application-specific integrated circuit (ASIC), and a printed circuit board with surface mounted components. Cell samples to be sequenced are processed through lysis, nucleic acid extraction, and other methods to extract long-chain DNA molecules. These DNA molecules are then subjected to DNA repair and adapter ligation. Subsequently, we mount flow cells in the socket of the CycloneSEQ sequencer and perform a chip self-check. After the self-check process, the system indicates whether the chip meets the quality criteria and the number of effective nanopores on each individual chip. After self-check process, we sequentially add the sequencing reagents and the library molecules to be sequenced into the micro-port of the chip, following a specific order. Then, we initiate the sequencing process through the software. Owing to the characteristics of nanopore single-molecule sequencing, as soon as the sequencing starts, the high-performance workstation paired with the sequencer can commence the base calling process. The CycloneSEQ sequencer is capable of supporting sequencing and base calling in real time simultaneously.

**Figure 4.**
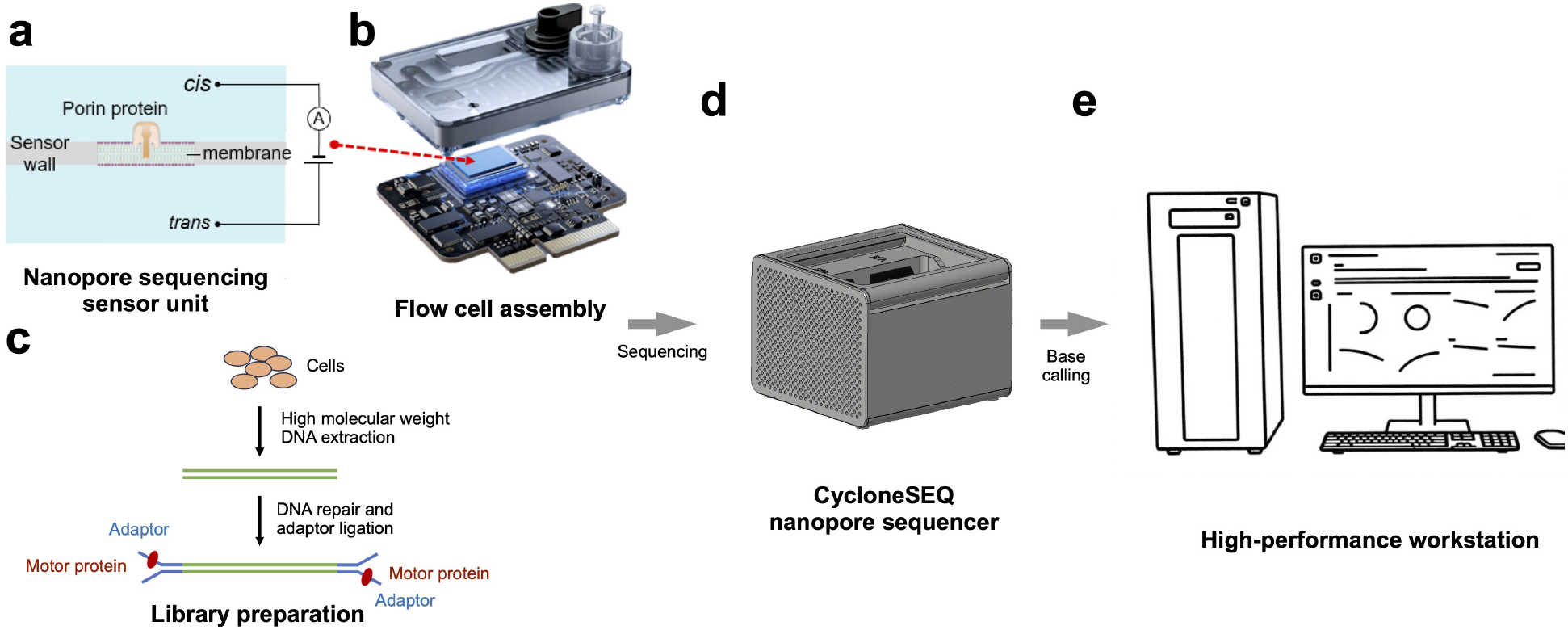
A single-molecule nanopore sequencing platform. **(a)** Schematic representation of a sensor unit constructed with an insulating membrane containing an inserted nanopore, along with *cis* and *trans* chambers and corresponding electrodes. **(b)** Exploded view diagram of a flow cell. **(c)** Library preparation process for nanopore sequencing. **(d)** CycloneSEQ nanopore sequencer. **(e)** High-performance workstation and operating software for the CycloneSEQ nanopore sequencing platform.

### 2.5 Error profile of CycloneSEQ

To systematically evaluate the performance of the CycloneSEQ platform, we generated whole-genome sequencing (WGS) data for the thoroughly characterized Genome in a Bottle (GIAB) consortium HG002 lymphoid cell line. Overall, the read lengths of the HG002 WGS data were distributed over a broad range from < 5 kb to > 50 kb, with a mean read length of 19.2 kb. The N50 value, defined as the length of the longest read that, together with longer reads, contain over 50% of all sequenced bases, was 33.6 kb (Fig. 5a). The mean bases quality values were predominantly in the range between 12 and 16, with a small cluster of short reads that have lower quality scores (Fig. 5b). The distribution of quality scores were fairly consistent in different relative positions of each read, and only dropped slighly near 5’ and 3’ ends (Fig. 5c).

**Figure 5.**
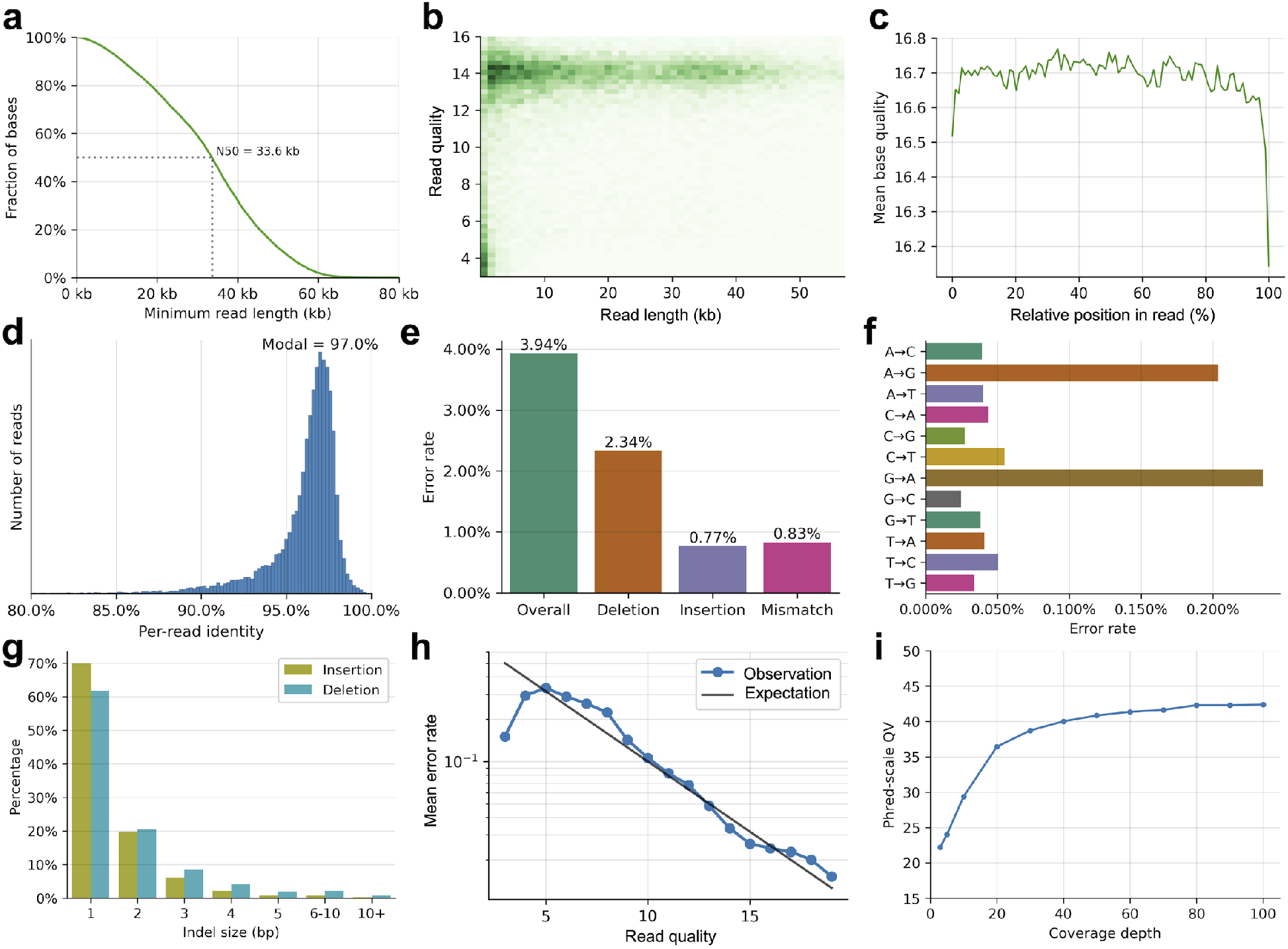
Whole-genome sequencing of the HG002 cell line. **(a)** The distribution of read lengths. The cumulative fraction of bases (y axis) contained in reads longer than a given threshold (x axis) is shown. Gray dotted lines denote N50, which represents the length of the longest read that, together with longer reads, contain over 50% of all sequenced bases. **b** Joint distribution of read lengths (x axis) and read quality scores (y axis). **(c)** Mean base quality (y axis) along different relative positions of the read (x axis). **(d)** Distribution of per-read identity. Per-read identity is calculated by aligning reads to the diploid HG002 v1.0.1 reference genome. **(e)** The overall error rate and the contributions of insertion, deletion and mismatch errors. **(f)** Contribution of each type of mismatch errors to the overall error rate. **(g)** Distribution of indel sizes for insertion and deletion errors. **(h)** Base quality calibration curve showing the relationship between per-read error rate (y axis) and per-read mean base quality (x axis). The blue line with dots represents observed results in CycloneSEQ data and the black line represents the expected error rates based theoretical calculations. **(i)** Consensus accuracy of CycloneSEQ reads (y axis) plotted against mean coverage depth (x axis). **(d)**-**(g)** are based on reads with mean base quality ≥ 10.

We aligned the sequencing reads to the diploid HG002 v1.0.1 reference genome in a haplotype-specific manner (see Methods), and analysed the frequency and types of sequencing errors from based on the alignment pattern. After removing reads with mean base quality scores below 10, which accounted for less than 10% of total bases, the accuracy of most reads ranged between 93% and 99%, with a modal value at 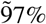 (Fig. 5d). The overall per-base error rate was estimated to be 3.94%, with deletions being the most frequent type of error, with an error rate contribution of 2.34%, followed by mismatches (0.83%) and insertions (0.77%) (Fig. 5e). For mismatch errors, A-to-G and G-to-A errors were significantly more common than other types of base substitutions, both of which had more than 0.2% error rate contributions (Fig. 5f). The enrichment of A-to-G and G-to-A errors was presumably due to the structural similarity between adenine and guanosine nucleotides, which lead to similar current signals [12]. Among the insertion and deletion errors, most errors affected either one or two bases, with only less than 10% insertion and deletion errors affecting three of more bases (Fig. 5g). Comparing per-read error rates estimated by reference alignment with reported mean base quality scores, we found that the reported quality scores was remarkably close to the actual sequencing accuracy (Fig. 5h). In contrast, similar systematic biases of quality scores often exist in other long-read sequencing platforms, which may affect downstream applications that rely on quality scores, such as variant calling [12]. We further analyzed the consensus accuracy of CycloneSEQ reads based on the *de novo* assemblies of the *E. coli* genome. The phred-scale quality value (QV) increased with coverage depth and reached 40 (i.e. error rate 0.01%) at 40 × coverage (Fig. 5i), confirming the possibility to acquire highly accurate consensus sequences using CycloneSEQ data alone.

### 2.6 Variant calling and *de novo* assembly of the HG002 genome

Variant calling and *de novo* assembly are among the most important applications of long-read sequencing in genomics research. For variant calling, long-read sequencing provides longer sequences that can resolve complex structural variants that are challenging for short-read sequencing. Longer reads also reduce ambiguity in read alignment, eliminating potential alignment errors in complex genomic regions. For *de novo* assembly, longer reads cover larger genomic regions in single reads, reducing computational complexity, and are more likely to span large repetitive elements, improving assembly contiguity. We observed that the coverage depth of CycloneSEQ reads were highly uniform across the human genome (except in the repeat-rich centromere regions prone to alignment errors), providing solid support for both variant calling and *de novo* assembly applications (Fig. 13).

Here, we generated variant calling and haplotype-resolved *de novo* assembly results for the HG002 genome using CycloneSEQ data. Variant calling was performed using our in-house bioinformatics tools LRAPmut and LRAPsv (see Methods) and compared against the Genome in a Bottle (GIAB) HG002 benchmark [13]. For single-nucleotide polymorphisms (SNPs), we achieved a precision of 0.992 and a recall of 0.990 at a sequencing depth of 30 × (Fig. 6). Small insertions and deletions (indels) present a challenge for variant calling due to their similarity to the predominant sequencing errors in CycloneSEQ data (Fig. 6). Utilizing variant imputation based on the 1000 Genomes reference panel (see Methods), we attained a precision of 0.955 and a recall of 0.890 at 30*×* coverage (Fig. 6). For structural variants (SVs), we observed that increased sequencing coverage had a more pronounced effect on improving precision and recall compared to SNPs and indels. Specifically, we achieved a precision of 0.948 and a recall of 0.954 at 40 × coverage (Fig. 6).

**Figure 6.**
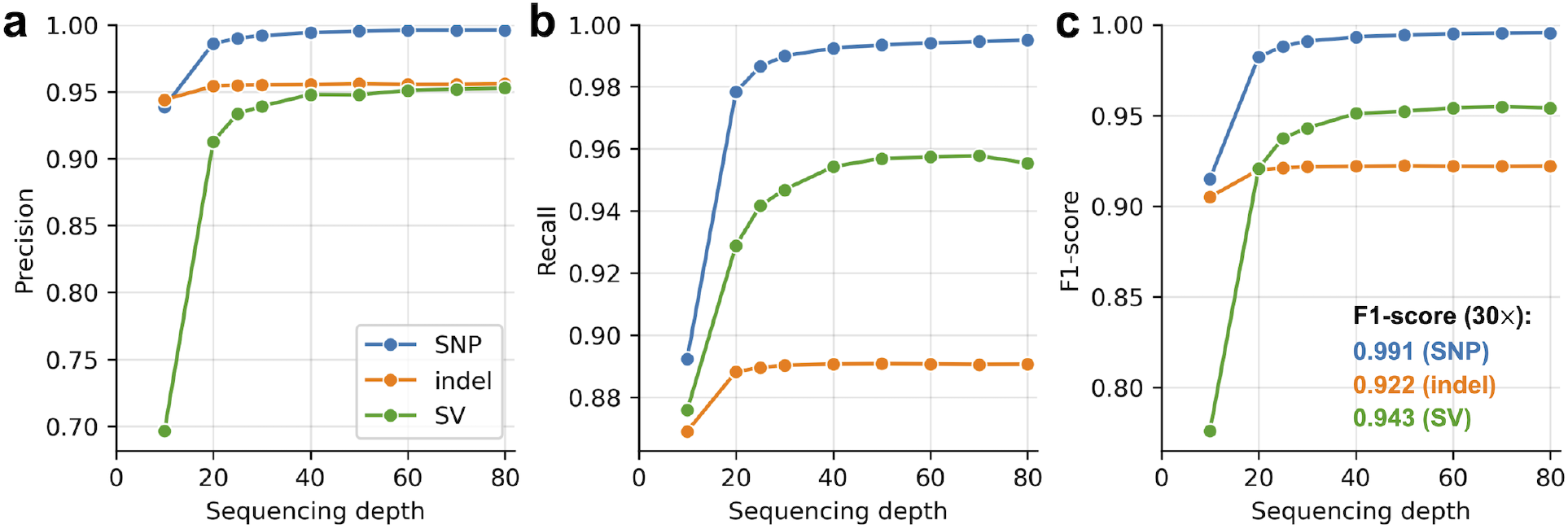
Variant calling of HG002. Precision **(a)**, recall **(b)** and F1-score **(c)** statistics of HG002 variant calling results based on CyclongSEQ data. The GIAB HG002 variant benchmark dataset was used as the ground truth.

The haplotype-resolved whole-genome *de novo* assembly for HG002 was generated using the Shasta assembler [14]. This assembly was evaluated against the Telomere-to-Telomere (T2T) consortium HG002 reference sequence (see Methods). We found that most chromosomes were assembled with a high level of completeness, with only the complex, repeat-rich centromere regions missing from the assembly (Fig. 7a). The short arms of the five acrocentric chromosomes—13, 14, 15, 21, and 22—were assembled with fragmented contigs due to the presence of satellite repeats and high sequence similarity among them. Other parts of the genome were mostly assembled with long, haplotype-resolved contigs, except the two sex chromosomes, likely due to the limited ability of the current Shasta implementagtion to handle the haplodity of X and Y chromosomes and the sequence homology between them. The NGA50 value of the assembly was 23.8 Mb (Fig. 7b), indicating that 50% of the genome was assembled with contigs of at least 23.8 Mb in length. The overall error rate of the assembly was estimated to be 0.12%, with deletions contributing the most to the overall error rate, followed by insertions and mismatch errors (Fig. 7c). Further developments on read lengths, accuracy and assembly methods will likely improve assembly contiguity and quality in the future.

**Figure 7.**
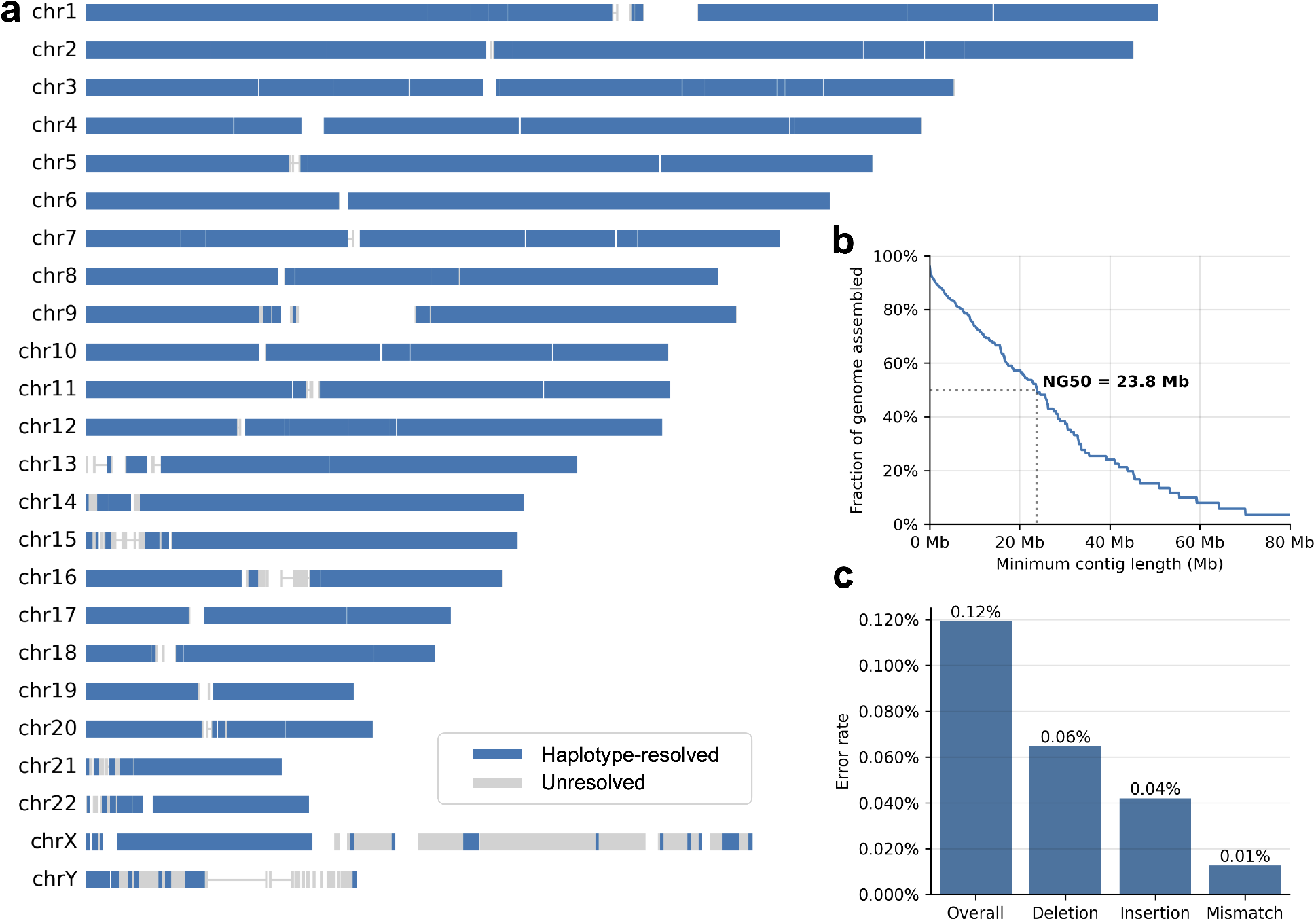
Haplotype-resolved whole-genome assembly for HG002. **(a)** Alignment of the assembled sequences to the T2T HG002 reference sequence. Each row represent a chromosome. Haplotype-resolved contigs and unresolved unitigs are shown in blue and gray, respectively. **(b)** Assembly contiguity represented as the fraction of genome assembled (y axis) for each minimum contig length (x axis). **(c)** Overall error rate of the assembly and error rates for deletion, insertion and mismatch errors.

### 2.7 Metagenome sequencing of mock sample

Metagenomic sequencing is an important application of nanopore sequencing. To evaluate the performance of CycloneSEQ in assembling microorganism genomes and estimating their relative abundance from a mixed sample, we generated 7.7 Gb sequencing data from the ZymoBIOMICS Gut Microbiome Standard mock metagenome sample, which contained a mixture of 17 microorganism species of predefined abundances. By alignment of CycloneSEQ reads to the corresponding reference genomes, we were able to accurately quantify the relative DNA abundance of 15 out of the 17 species in the sample based on sequencing depths, including both high-GC and low-GC species (Fig. 8a). Only two of the least abundant species could not be identified from sequencing data: *Enterococcus faecalis* (abundance 0.001%) and *Clostridium perfringens* (abundance 0.0001%).

**Figure 8.**
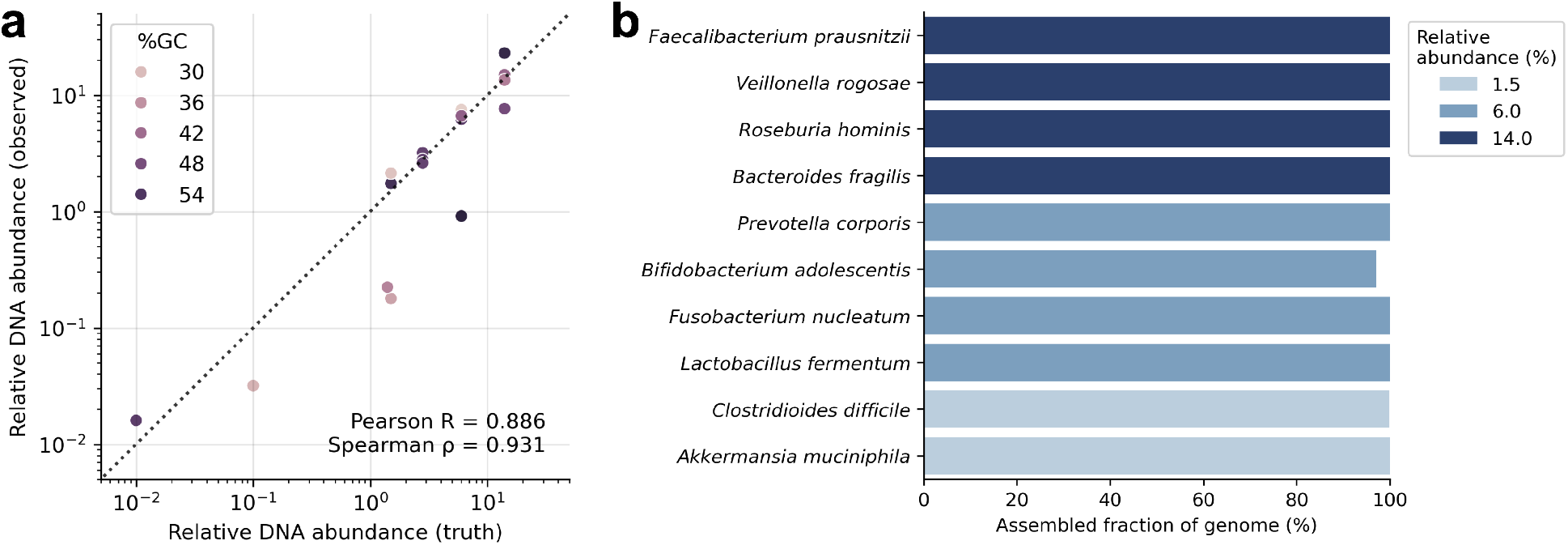
Metagenome sequencing and methylation prediction. **(a)** Correlation between expected (x axis) and observed (y axis) relative DNA abundance in the mock metagenome sample. Each dot represents a microorganism strain. Dot colors represent the GC content of the genome of the assayed strains. **(b)** Assembled genome fraction of ten species in the mock metagenome sample. Colors represent the relative DNA abundances.

In addition to reference-based quantification, we performed *de novo* assembly using the Flye assembler [15] based on CycloneSEQ data. Among the 17 species in the sample, ten species had relative DNA and genome copy abundances above 1%, all of which were assembled with high levels of genome completeness, with nine out of ten genomes successfully circularized (Fig. 8b and Table 1). Based on the assembled genomes, we were also able to perform accurate quantification of the copy numbers and sequence lengths of 16S, 5S and 23S rRNA of these ten species (Tables 2, 3, and 4).

**Table 1:** The relative abundance, genome size, and assembled fraction of different species in the mock metagenome sample. Species with relative genomic DNA or genome copy abundances below 1% are excluded.

| Species | Relative abundance (%) | Genome size (Mb) | Assembled fraction (%) |
| --- | --- | --- | --- |
| *Faecalibacterium prausnitzii* | 14 | 2.914 | 100 |
| *Veillonella rogosae* | 14 | 2.158 | 100 |
| *Roseburia hominis* | 14 | 3.463 | 100 |
| *Bacteroides fragilis* | 14 | 5.167 | 99.988 |
| *Prevotella corporis* | 6 | 2.947 | 99.977 |
| *Bifidobacterium adolescentis* | 6 | 2.090 | 97.067 |
| *Fusobacterium nucleatum* | 6 | 2.448 | 99.96 |
| *Lactobacillus fermentum* | 6 | 1.905 | 99.997 |
| *Clostridioides difficile* | 1.5 | 4.209 | 99.894 |
| *Akkermansia muciniphila* | 1.5 | 2.851 | 100 |

**Table 2:**
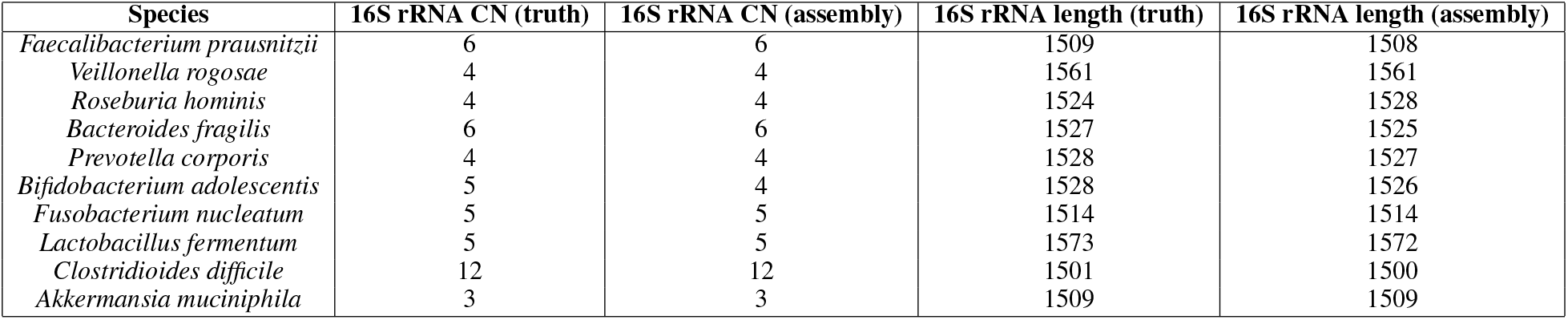
Statistics of 16S rRNA copy numbers (CN) and lengths of different species in the mock metagenome sample. Species with relative genomic DNA or genome copy abundances below 1% are excluded.

**Table 3:** Statistics of 5S rRNA copy numbers (CN) and lengths of different species in the mock metagenome sample. Species with relative genomic DNA or genome copy abundances below 1% are excluded.

| Species | 5S rRNA CN (truth) | 5S rRNA CN (assembly) | 5S rRNA length (truth) | 5S rRNA length (assembly) |
| --- | --- | --- | --- | --- |
| *Akkermansia muciniphila* | 3 | 3 | 108 | 108 |
| *Bacteroides fragilis* | 6 | 6 | 106 | 106 |
| *Bifidobacterium adolescentis* | 6 | 5 | 109 | 109 |
| *Clostridioides difficile* | 11 | 11 | 105 | 106 |
| *Faecalibacterium prausnitzii* | 6 | 6 | 112 | 112 |
| *Fusobacterium nucleatum* | 5 | 5 | 107 | 107 |
| *Lactobacillus fermentum* | 5 | 5 | 112 | 112 |
| *Prevotella corporis* | 4 | 4 | 109 | 109 |
| *Roseburia hominis* | 4 | 4 | 100 | 100 |
| *Veillonella rogosae* | 4 | 4 | 110 | 110 |

**Table 4:** Statistics of 23S rRNA copy numbers (CN) and lengths of different species in the mock metagenome sample. Species with relative genomic DNA or genome copy abundances below 1% are excluded.

| Species | 23S rRNA CN (truth) | 23S rRNA CN (assembly) | 23S rRNA length (truth) | 23S rRNA length (assembly) |
| --- | --- | --- | --- | --- |
| Akkermansia muciniphila | 3 | 3 | 2830 | 2830 |
| Bacteroides fragilis | 6 | 6 | 2880 | 2880 |
| Bifidobacterium adolescentis | 4 | 5 | 3040 | 3049 |
| *Clostridioides difficile* | 12 | 13 | 2893 | 2894 |
| Faecalibacterium prausnitzii | 6 | 6 | 2831 | 2831 |
| Fusobacterium nucleatum | 5 | 5 | 2892 | 2891 |
| *Lactobacillus fermentum* | 5 | 5 | 2917 | 2917 |
| *Prevotella corporis* | 4 | 4 | 2892 | 2894 |
| Roseburia hominis | 4 | 4 | 2884 | 2885 |
| Veillonella rogosae | 4 | 4 | 2927 | 2928 |

### 2.8 Single-cell RNA sequencing of GM12878 cell line

Long-read sequencing in single-cell RNA sequencing (scRNA-seq) offers significant advantages, including the ability to capture full-length transcripts, which provides a more comprehensive view of isoform diversity, alternative splicing, and gene fusion events. This detailed transcriptome profiling enhances the understanding of cellular heterogeneity and complex gene regulatory networks. Here, we present preliminary results on applying the CycloneSEQ platform in scRNA-seq using mRNA from the GM12878 cell line, and compare the results with scRNA-seq data generated from the same cDNA libraries using BGI DNBSEQ short-read sequencing. As most transcripts were less than 10 kb in length, it was possible to obtain full-length coverage from single CycloneSEQ long reads. We found that the mean coverage depth was slightly higher near the 3’ end of each gene and lower near the 5’ end of each gene (Fig. 9a), likely due to incomplete reverse transcription and/or degregation of mRNA near 5’ ends. Despite this, the overall coverage depths were highly uniform from the 5’ to 3’ end of each transcript (Fig. 9a), providing solid support for the discovery of potential novel isoforms. The total number of genes detected in each cell by CycloneSEQ ranged between 300 and 4,000 and showed a strong linear correlation (*R*^2^ = 0.95) with that of DNBSEQ data. Gene expression quantification results from CycloneSEQ were also highly consistent (*R*^2^ = 0.93) with those of DNBSEQ short-read sequencing, suggesting that our CycloneSEQ platform was capable of accurate transcript quantification in single-cell sequencing.

**Figure 9.**
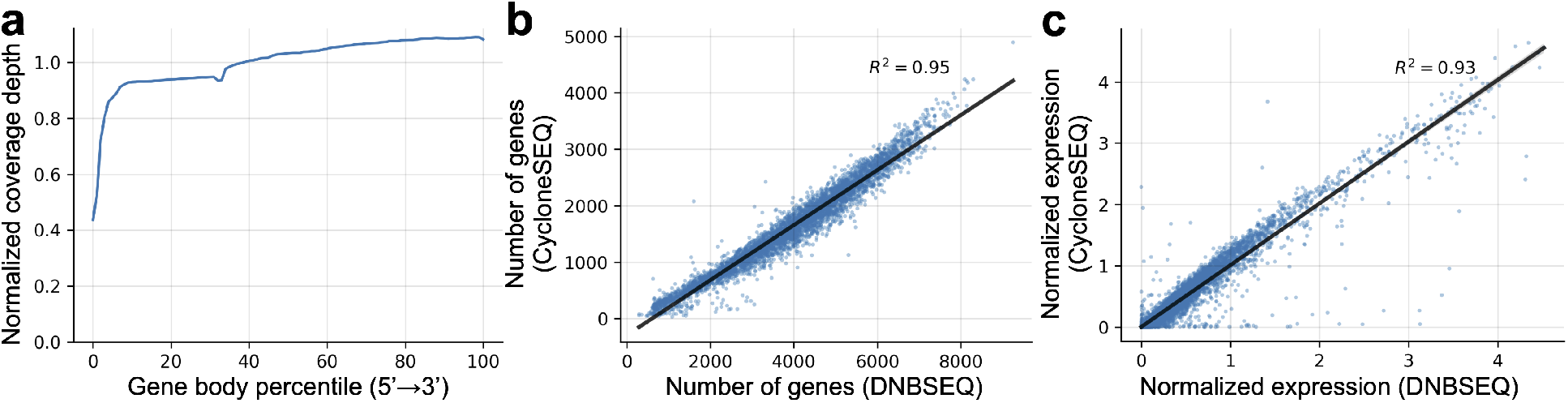
Single-cell RNA sequencing of GM12878 cell line. **(a)** Normalized coverage depths (y axis) along different relative positions of the gene body (x axis). Transcripts shorter than 100 bp are not included. **(b)** Correlation between the number of detected gene in each cell by DNBSEQ (x axis) and that by CycloneSEQ (y axis) platforms. Each dot represents a cell. *R* represents the Pearson regression coefficient. **(c)** Correlation between pseudobulk gene expression levels measured by DNBSEQ (x axis) and that by CycloneSEQ (y axis) platforms. Each dot represents a gene detected by both platforms. *R* represents the Pearson regression coefficient.

**Figure 10.**
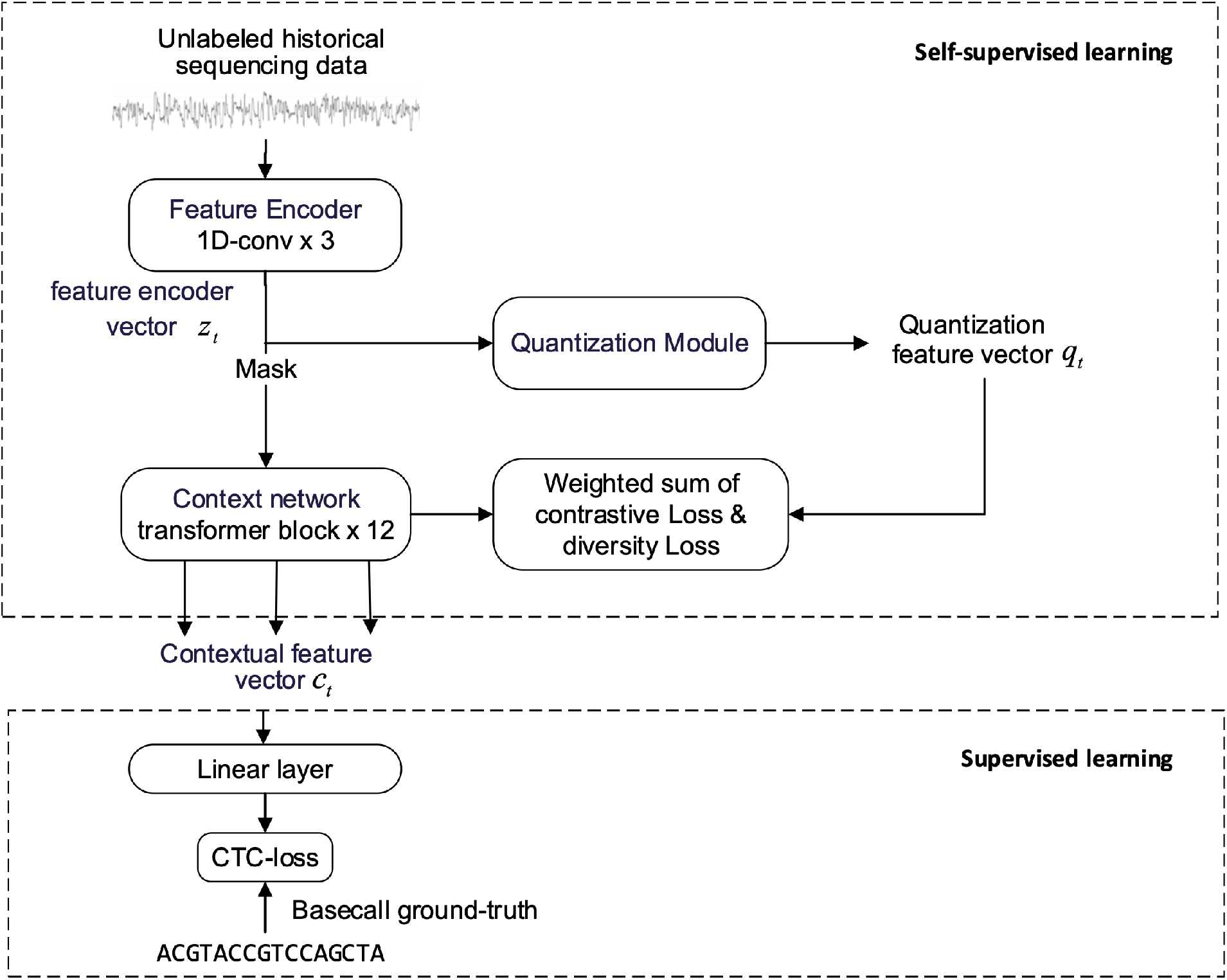
Architecture of the pre-training model for base-calling

## 3 Discussions

Nanopore sequencing has revolutionized genomics by enabling the real-time analysis of nucleic acids without the need for amplification or chemical labeling. Despite its transformative impact, several limitations, such as relatively high error rate and low throughput, still hinder its broader adoption and effectiveness. In this study, we developed several novel components of nanopore sequencing technology, including motor and pore proteins, chip design, and basecalling algorithms. We show that our new sequencing platform, CycloneSEQ, is capable of generating high-throughput long-read sequencing data useful for downstream applications.

One significant limitation of current nanopore sequencing technology is its accuracy. Although improvements have been made, the error rate remains higher compared to short-read sequencing technologies. This issue can be addressed by screening for novel motor and pore proteins and developing more advanced basecalling algorithms that could enhance the precision and speed of nanopore sequencing [16]. Future development and integration of non-protein pores, such as synthetic nanopores, may provide more stable and consistent results, reducing the error rates and increasing the reliability of the technology [17]. Another area of improvement lies in the membranes used for nanopore sequencing. Novel membranes with enhanced stability and reduced noise could significantly improve the quality of the sequencing data [18]. Advances in materials science could lead to the development of membranes that are more resilient to the harsh conditions often encountered during sequencing, thereby extending their lifespan and efficiency.

Sequencing of modified DNA and RNA bases remains a challenge for nanopore technology. Modified bases play crucial roles in various biological processes, and accurate detection is essential for understanding epigenetics and other regulatory mechanisms. Enhancing the capability of nanopore sequencing to accurately read modified bases would be a significant breakthrough, potentially achieved through the use of advanced bioinformatics algorithms and improved pore chemistry [19].

The potential of nanopore technology extends beyond nucleic acids to the sequencing of amino acids, peptides, and proteins. Protein sequencing using nanopores could revolutionize proteomics, enabling the direct analysis of proteins and their post-translational modifications. Although still in its infancy, this application holds promise for significant advancements in understanding protein structure and function [20].

Clinical applications of nanopore sequencing are vast, ranging from rapid pathogen identification to point-of-care testing. However, the high costs and limited throughput of current systems restrict their widespread use in clinical settings. Reducing costs and improving throughput are critical for the adoption of nanopore sequencing in routine clinical practice [8][21]. Innovations in sequencing chemistry, automation, and data analysis could make nanopore sequencing more accessible and practical for clinical diagnostics. Additionally, nanopore sequencing holds great potential for population cohort studies, providing insights into genetic diversity and disease susceptibility on a large scale. The ability to sequence entire genomes quickly and cost-effectively, and accurately characterize structure variation could transform epidemiological studies and public health initiatives. Continued advancements in reducing costs and increasing throughput are essential to fully realize the potential of nanopore sequencing in both clinical and population cohort applications.

## 4 Methods

### 4.1 Development of Motor Proteins and Pore Proteins

All protein sequences were derived from a deep-sea metagenomic database, and all mutant designs were based on AlphaFold3 [9] structure predictions. The proteins were overexpressed in BL21(DE3) or similar strains, followed by purification using affinity chromatography, ion exchange chromatography, and size-exclusion chromatography. DNA libraries were prepared from the motor protein BCH-X mutants, Y-shaped adaptors and input DNA. After embedding the pore protein BCP-Y mutants into membranes, sequencing buffer and the test libraries were added, and then sequencing was performed at 0.18V and 30°C to collect current signals. The sequencing speeds were obtained by dividing the length of the specific sequence DNA by its translocation time. The open pore currents of BCP-Y mutants were recorded at different voltages (0V, 0.02V, 0.04V, 0.10V, 0.14V, and 0.18V).

### 4.2 Training and validation of basecalling models

#### 4.2.1 Model architecture

Our model consists of three key components (Fig. 11): (1) Feature Encoder: A multi-layer 1D convolution network processes raw signals through feature encoding, capturing relevant information from the sequencing data. It includes multiple blocks, each with 1D convolution and GELU activation for downsampling and extracting local patterns; (2) Quantization Module: The feature encoder output is discretized using product quantization into a finite representation space, enhancing the model’s self-supervised learning capability. (3) Mask and Context Networks: The feature encoder outputs are processed by a mask module before being fed into the context network, composed of multiple transformer layers. This network approximates relative positional encoding and enhances contextual understanding.

#### 4.2.2 Improved definition of the Contrastive Loss

The objective function of self-supervised pre-training comprises two components: Contrastive Loss and Diversity Loss. These components optimize different aspects of the training process to improve model performance. Contrastive Loss measures the contextual network’s ability to predict future outputs. The Transformer’s output at time *t* (*c*_*t*_) should match the quantization module’s output at time *t* (*q*_*t*_). Additionally, *c*_*t*_ should differ from outputs at *k* other randomly selected positions (distractors) in the sequence. The original Contrastive Loss is defined as:

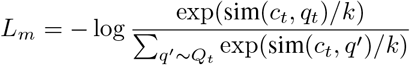

Where 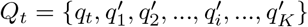, sim() is a predefined similarity function, 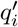 is the i-th interference vector, and *k* is a predefined temperature parameter.

We identified an issue where positive and negative samples could map to the same quantized vector, undermining performance. Assuming *q*_*t*_ is the positive sample’s quantized vector at time *t*, the corresponding negative sample quantized vectors are divided into *q*^*′*^, not equal to *q*_*t*_, and *p*^*′*^, equal to *q*_*t*_:

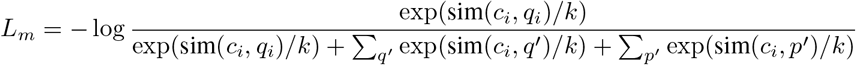

Negative samples mapping to the same quantized vector as positive samples undermine contrastive loss. In wav2vec 2.0, such negatives do not contribute to the contrastive loss, making the last denominator term zero. This issue halts the training process when contrastive loss becomes zero, preventing gradient updates.

We introduced a penalty count(*p*^*′*^) *· l* for such cases, with *l* as the penalty coefficient (*l* = 0.01):

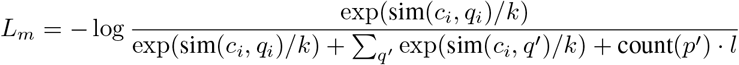

Our method applies penalties to both Contrastive Loss and Diversity Loss, ensuring negative samples contribute meaningfully. This improvement reduces instances where contrastive loss reaches zero, enhancing training robustness and effectiveness.

#### 4.2.3 Pre-training and fine-tuning

The pre-trained data originated from the BGI Cyclone nanopore sequencing platform, including sequences from various species: 80% human genome, 20% rice, *Saccharomyces cerevisiae*, and *Bacillus subtilis*. Data was sampled at 5 kHz, filtering abnormal signals, and divided into chunks of 5000 signal points, creating 300 million pre-training chunks. We used a 15% probability mask, mask length of 5. The feature encoder employed a 3-layer 1D convolution with a stride of (1,1,5), kernel size of (5, 5, 19), and dimension of (4, 16, 768). The context network utilized a 10-layer transformer with 512 hidden units and 8 heads. Training on 64 NVIDIA A100-PCIE-40GB GPUs for 3.5 days, batch size 1024, using Adam optimizer with learning rate 0.005 and linear decay. *α* = 0.1 for diversity loss, *K* = 100 for negative examples, *G* = 2 and *V* = 320 for the quantization module. Fig. 12 shows training and validation loss over iterations.

**Figure 11.**
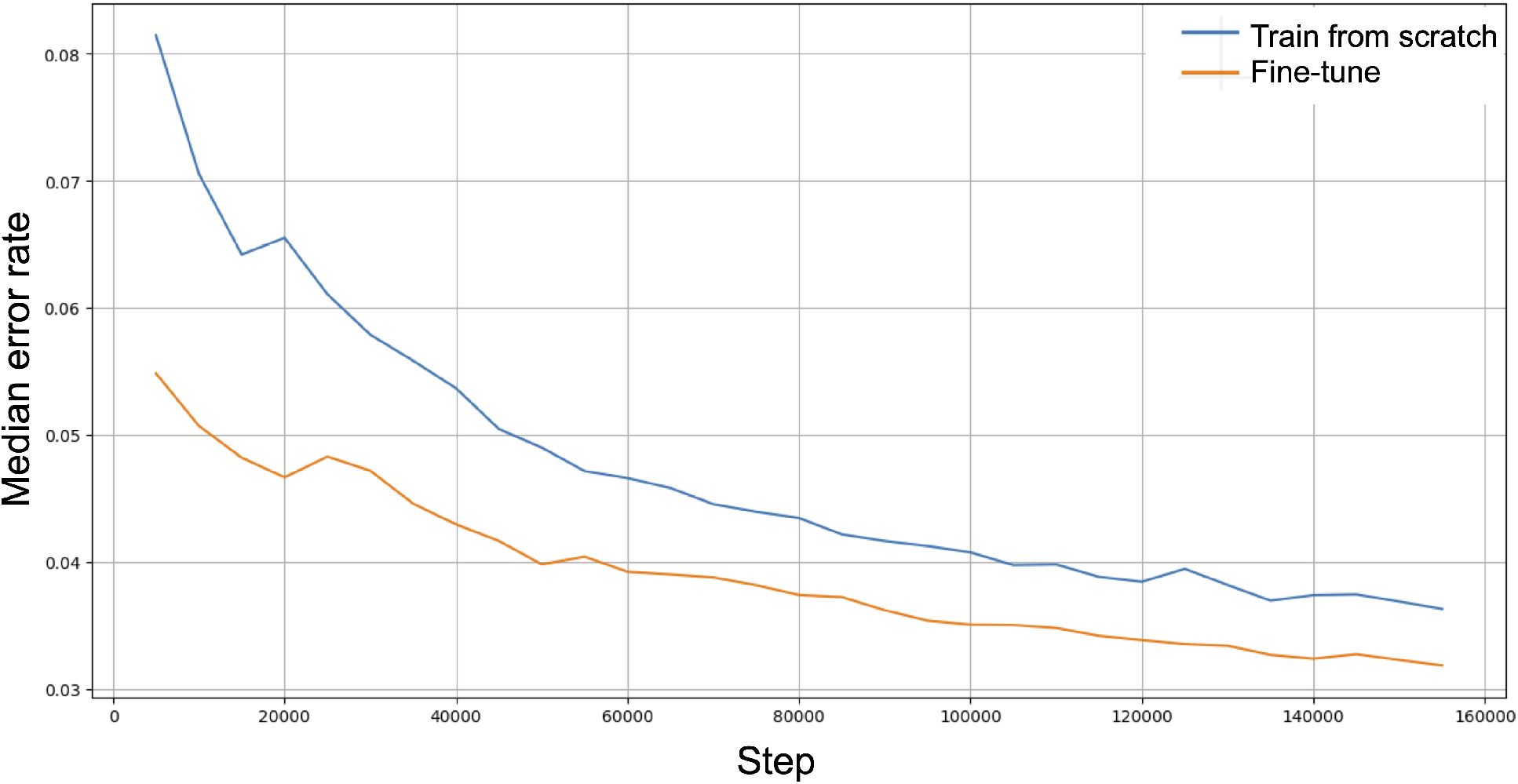
Performance of fine-tuned basecalling model

**Figure 12.**
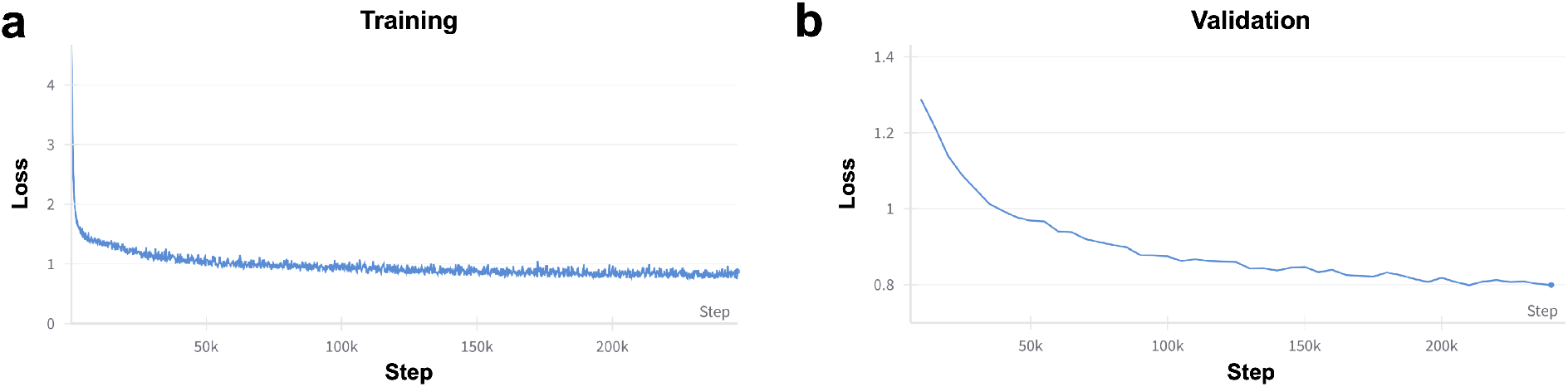
Training **(a)** and validation **(b)** loss in the pre-train stage of the basecalling model.

**Figure 13.**
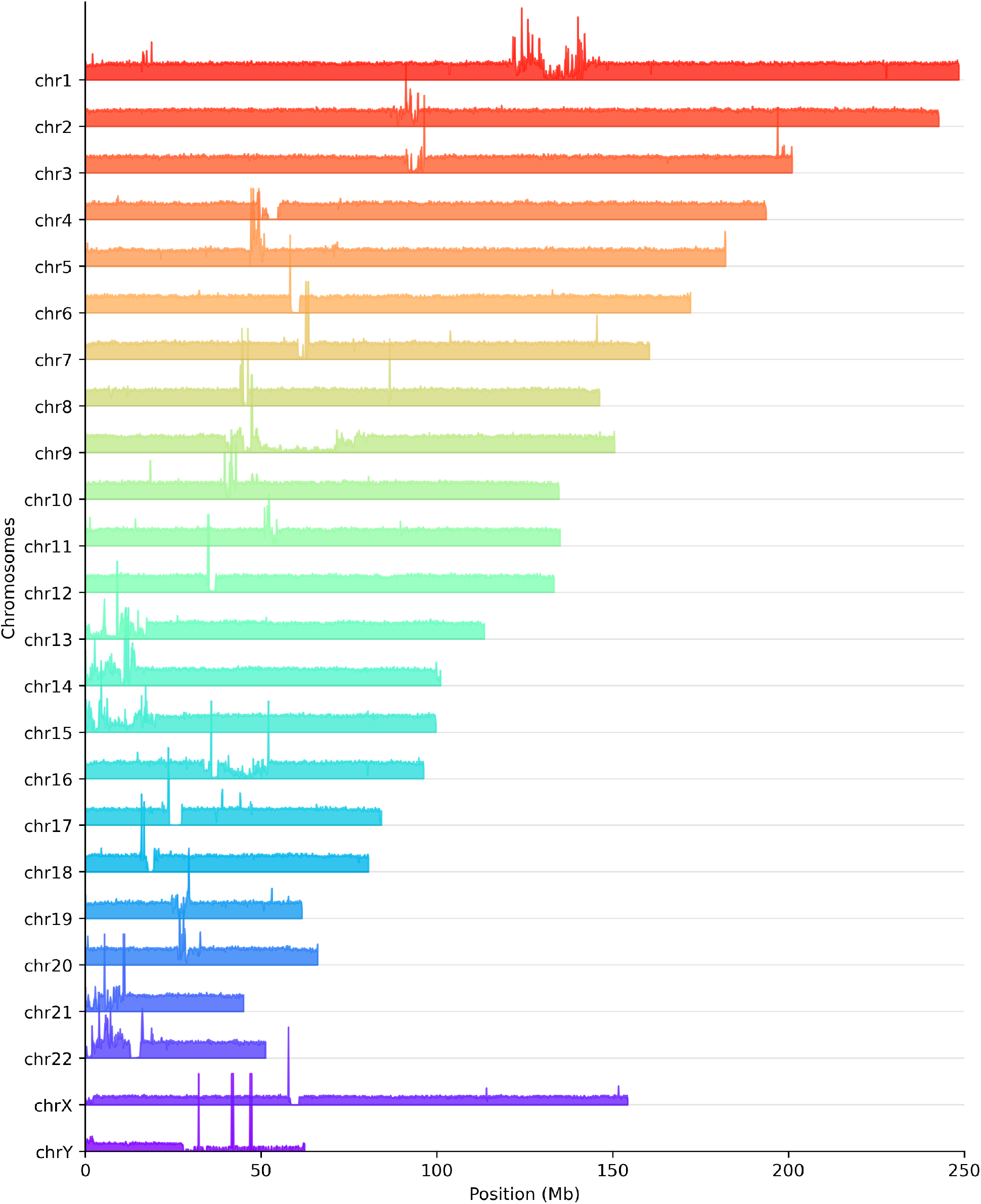
Coverage depth of CycloneSEQ data in the HG002 genome.

Our pre-training utilized a dual loss function: Contrastive Loss to gauge the context network’s predictive capability, and Diversity Loss to enhance the quantization codebooks’ expressiveness. We identified a flaw in wav2vec 2.0’s handling of contrastive loss, where positive and negative samples could map to the same quantized vector. Our improvement penalizes such occurrences, reducing cases where contrastive loss approaches zero and enhancing training effectiveness (see Methods). After pre-training, the model was fine-tuned for base identification. A linear layer maps the output to categories representing the four nucleobases and a placeholder. We used CTC-Loss for optimization. The fine-tuned model showed marked improvements in error rates and faster convergence compared to models initialized randomly (Fig. 10), highlighting the effectiveness of leveraging pre-trained weights, especially with limited labeled data. Experiments on human and other species data indicated that pre-training enables the model to generalize across species, evidenced by superior performance on species not covered during pre-training. Additionally, non-masked fine-tuning yielded better results, contrary to wav2vec 2.0’s findings [11], suggesting task-specific differences in optimal training strategies.

Fine-tuning used data from the BGI Cyclone sequencing platform. An existing basecaller (Bonito) predicted base sequences, compared with a standard library, selecting those with coverage >95% as labels. Post pre-training, the model was fine-tuned on 40 million annotated human samples, with 10,000 evaluation samples, for 1.3 days on 16 NVIDIA A100-PCIE-40GB GPUs, batch size 256. Using Adam optimizer with a warm-up over 200 steps followed by linear decay, we experimented with learning rates of 0.0005 and 0.001, reporting the best results. Models were evaluated using median error rate.

### 4.3 Nanopore local chemistry (NLC) sequencing

We established a model in COMSOL to simulate and analyze the concentration distribution of Mg^2+^ in the *cis* and *trans* chambers on both sides of the nanopore. The system model parameters were set as follows: the nanopore opening diameter was 2 nm, the length was set to 10 nm, and the width and height of both the *cis* and *trans* chambers were set to 100 nm and 52.5 nm, respectively. The *cis* chamber was filled with a 0.5 M KCl aqueous solution, and the *trans* chamber was filled with a 0.5 M KCl and 20 mM MgCl_2_ aqueous solution. In this model, we considered two main transport mechanisms: ion diffusion and ion electrophoresis. Given that the nanopore interior was assumed to be neutral without any charge settings, the effect of electroosmotic flow was neglected. Different trans-membrane potentials were applied to the system, and the steady-state Mg^2+^ concentration gradient distribution was calculated via COMSOL.

As shown in Fig. 2c, the lower chamber was subjected to a boundary condition potential of 0 V, and the upper chamber was grounded at 0 V, resulting in no applied potential difference across the nanopore. Under this condition, there was no significant ion electrophoresis behavior guided by an electric field. The concentration gradient distribution of Mg^2+^ was primarily determined by free diffusion of ions. As shown in Fig. 2d, when the lower chamber was subjected to a boundary condition potential of 0.18 V and the upper chamber was grounded at 0 V, an applied potential difference of approximately 0.18 V was established across the nanopore.

### 4.4 Nanopore sensor chip

The nanopore sensor chip was fabricated using standard semiconductor manufacturing techniques, including photolithography, thin film deposition, and patterning. The confocal image was captured using a Nikon C2+ microscope. The membrane solution was mixed with 2 µM BODIPY PL fluorescent dye, which has an excitation peak at 502 nm and an emission peak at 511 nm. The electrolyte in the microwell was stained with 0.02% wt Sulforhodamine B, which has an excitation peak at 559 nm and an emission peak at 577 nm.

### 4.5 DNA Extraction and Library Preparation

High molecular weight DNA extraction and optional length sorting were performed on the sample to obtain DNA suitable for long-read nanopore sequencing. The quality of the extracted DNA was assessed by measuring the A260/A280 ratio, which was maintained between 1.8 and 2.0. End repair reagents were used to repair the ends of the DNA fragments and add deoxyadenosine (dA) tails, which facilitated subsequent adaptor ligation. Oligos were annealed in TE buffer to form Y-shaped adaptors. The sequencing library was prepared from motor protein BCH-X, Y-shaped adaptors and input DNA.

### 4.6 Sequencing error analyses

For sequencing error analyses, low-speed sequencing mode was used to generate the HG002 whole genome sequencing data on the CycloneSEQ platform. We first randomly sampled 20,000 reads from the sequencing data, which were aligned to both haplotypes of the diploid reference genome HG002 v1.0.1 using Minimap2 [22]. Insertion, deletion and mismatch errors were identified from the CIGAR string of the resulting alignments. Error rates were calculated by dividing the total length of errors by the total alignment length. To compute read quality for each read, we first converted Phred-scale base quality values to error rates, and then calculated the average error rate of all bases in each read. The resulting average error rate was finally converted back to Phred-scale to represent the read quality of each read. Visualization was performed using Matplotlib [23] and Seaborn [24] in Python.

### 4.7 Variant calling

We used minimap2 version 2.24-r1164-dirty to align reads to a reference genome. For small variant calling, we utilized LRAPmut version v1.0 (https://github.com/Roick-Leo/LRAPmut) with CycloneSEQ data, and for structural variant calling, we employed LRAPsv version v1.0 (https://github.com/Roick-Leo/LRAPsv). Haplotype imputation based on 1000 genomes reference panel was applied to improve the performance of indel variant calling. HG002 data was removed from the reference panel prior to imputation. To benchmark the variant calls, we assessed the small variant calls against the GIAB truth set using hap.py v0.3.15 (). Additionally, we used Truvari v4.2.2 (ref [25]) to produce performance metrics by comparing the predicted structural variants with the benchmark SVs.

### 4.8 De novo assembly of the HG002 genome

The haplotype-resolved \textit{de novo} assembly of the HG002 genome was generated using Shasta v0.11.1 [14], based on CycloneSEQ reads at approximately 80× genomic coverage. The following command was used:

~~~
shasta-Linux-0.11.1 \
  --input <INPUT_FASTQ> \
  --assemblyDirectory <OUTPUT_FOLDER> \
  --config Nanopore-Phased-May2022 \
  --threads 16 \
  --memoryMode filesystem \
  --memoryBacking disk
~~~

Assembly evaluation was performed by aligning the assembly to the T2T HG002 reference sequence (https://github.com/marbl/HG002). Each assembly contig was separately aligned to the paternal and maternal haplotypes of the HG002 reference sequence. The alignment with the largest number of matched bases was selected for evaluation. Deletion, insertion, and mismatch error rates were calculated based on the CIGAR string of the resulting alignment.

### 4.9 Assembly and quantification of a mock metagenome sample

The ZymoBIOMICS Gut Microbiome Standard (Catalog No. D6331) comprises 18 bacterial strains, 2 fungal strains, and 1 archaeal strain, with a theoretical genomic DNA abundance ranging from 0.0001% to 14%. Reference genomes, along with 16S and 18S rRNA genes, are accessible online [26]. DNA was extracted from the mock samples using the MGIEasy Stool Microbiome DNA Extraction Kit according to the manufacturer’s protocols.

Chopper 0.6.0 (https://github.com/wdecoster/chopper) was utilized to filter out reads with a quality score lower than Q10 and a length shorter than 1,000 base pairs using the following parameters: `-q 10 –minlength 1000`. Then, the reads were assembled using the Flye 2.8.3-b1695 using the following parameters: `–meta –nano-raw`. Semibin2 2.1.0 was applied to generate bins for the mock community using the following parameters: (`single_easy_bin –environment global –sequencing-type long_read`). Assembly quality was assessed using Quast v5.0.2. Then, the rRNA genes were predicted using barrnap 0.9 using the following parameters: `–kingdom -reject 0.01 -evalue 1e-3`. tRNA was predicted by tRNAscan-SE 2.0.12 using the following parameters: `-B`. For quantification and evaluation, plasmid sequences were removed from the reference sequences. Relative DNA abundances of each strain was estimated by aligning reads to the combined reference sequences of all strains, and summarising the total number of aligned bases for each strain.

### 4.10 Single-cell RNA sequencing of GM12878 cell line

For DNBSEQ sequencing, sample preparation, cell isolation, mRNA extraction, reverse transcription, library preparation, sequencing were performed according to the manufacturer’s guidelines of the DNASEQ C4 platform. For CycloneSEQ sequencing, the same procedure was followed with the addition of PCR amplification of cDNA libraries. The amplified libraries was sequenced on two CycloneSEQ chips. scRNAseq data analyses was performed using an in-house pipeline based on GENCODE GRCh38 transcript annotations.

## 5 Acknowledgements

This work was supported by “Pioneer” and “Leading Goose” R&D Program of Zhejiang (2024C03004), Shenzhen Science and Technology Program (KQTD20221101093603011), and National Key Research and Development Program of China (2022YFF1202103).

## 7 Supplementary information

### 7.1 Supplementary figures

### 7.2 Supplementary tables

